# Secondary analysis of transcriptomes of SARS-CoV-2 infection models to characterize COVID-19

**DOI:** 10.1101/2020.08.27.270835

**Authors:** Sudhir Ghandikota, Mihika Sharma, Anil G. Jegga

## Abstract

Knowledge about the molecular mechanisms driving COVID-19 pathophysiology and outcomes is still limited. To learn more about COVID-19 pathophysiology we performed secondary analyses of transcriptomic data from two *in vitro* (Calu-3 and Vero E6 cells) and one *in vivo* (Ad5-hACE2-sensitized mice) models of SARS-CoV-2 infection. We found 1467 conserved differentially expressed host genes (differentially expressed in at least two of the three model system transcriptomes compared) in SARS-CoV-2 infection. To find potential genetic factors associated with COVID-19, we analyzed these conserved differentially expressed genes using known human genotype-phenotype associations. Genome-wide association study enrichment analysis showed evidence of enrichment for GWA loci associated with platelet functions, blood pressure, body mass index, respiratory functions, and neurodegenerative and neuropsychiatric diseases, among others. Since human protein complexes are known to be directly related to viral infection, we combined and analyzed the conserved transcriptomic signature with SARS-CoV-2-host protein-protein interaction data and found more than 150 gene clusters. Of these, 29 clusters (with 5 or more genes in each cluster) had at least one gene encoding protein that interacts with SARS-CoV-2 proteome. These clusters were enriched for different cell types in lung including epithelial, endothelial, and immune cell types suggesting their pathophysiological relevancy to COVID-19. Finally, pathway analysis on the conserved differentially expressed genes and gene clusters showed alterations in several pathways and biological processes that could enable in understanding or hypothesizing molecular signatures inducing pathophysiological changes, risks, or sequelae of COVID-19.

## INTRODUCTION

Of the seven coronaviruses known to infect human beings, four (229E, HKU1, NL63, and OC43) are associated with several respiratory conditions ranging from common cold to bronchiolitis and pneumonia but rarely result in any mortality. The other three, namely, MERS-CoV, SARS-CoV, and the recent SARS-CoV-2, have different degrees of lethality. The SARS-CoV outbreak in 2002-2003 led to about 10 % mortality among the infected [1, 2]. The MERS-CoV, on the other hand killed about 23% of the infected people between 2012-2019 [3]. SARS-CoV-2 [4] which emerged in December 2019 in Wuhan of China, has already killed several more people than the SARS-CoV and MERS-CoV outbreaks combined. Since the initial report of coronavirus disease 2019 (COVID-19) in China on 30th December 2019 [5], as of August 26, 2020, there have been more than 24 million confirmed infections and >820 thousand deaths reported worldwide from 216 different countries (World Health Organization, 2020) including ~5.7 million confirmed infections and ~177,000 deaths in the USA. The COVID-19 pandemic dealt a devastating blow to global financial and social fabric and the impact on human health is expected to loom over several years to come if not decades. The limited and emerging stages of data and information surrounding this disease, and the necessity to find effective interventions (vaccines, small molecules, etc.), supplies a strong rationale for secondary analysis of existing data collected from different models and studies.

In the past 8 months, studies on COVID-19 included observational studies (e.g., clinical cohorts, epidemiological investigations and forecasts), in-silico genomic and structural analyses, and studies addressing basic virologic questions of SARS-CoV-2 (host cell tropism, viral replication kinetics, and cell damage profiles). Few studies have focused on the COVID-19 sequelae in surviving or recovered patients, genetic predisposition, and long-term effects if any in infected but asymptomatic individuals. Leveraging the available repository of datasets and information, even if they were not designed specifically to study COVID-19, can supply a jump start to discover different sides of this disease. Our study is based on the premise that combining information from multiple layers of data may result in new biologically interpretable associations in several ways. Indeed, some of the noteworthy discoveries surrounding SARS-CoV-2 are direct offshoots of secondary data analysis using available omics data generated in pre-COVID-19 times. For instance, in two seminal studies, Sungnak et al. [6] and Ziegler et al. [7] surveyed expression of SARS-CoV-2 entry-associated genes (*ACE2, TMPRSS2, CTSB*, and *CTSL*) in single-cell RNA-sequencing (scRNA-seq) data from multiple tissues from healthy human donors generated by the Human Cell Atlas consortium and other published and unpublished studies to investigate SARS-CoV-2 tropism. Likewise, there have been several other studies assessing the expression patterns of SARS-CoV-2 [8–11] and other coronavirus entry proteins [12] in barrier tissues. In yet another study, Genotype Tissue Expression (GTEx) database [13] was used to compare the genomic characteristics of ACE2 among different populations and systematically analyze coding-region variants in ACE2 and the eQTL variants that may affect the expression of ACE2 [14]. Characterizing COVID-19 pathophysiology and understanding the molecular effects of SARS-CoV-2 on human proteins is critical to discover, guide, and prioritize therapeutic strategies. In the current study, we leverage transcriptomic data from 3 model systems (two in vitro and one in vivo) of SARS-CoV-2 infection, SARS-CoV-2 viral-host protein interaction data, and analyze them jointly with non-COVID-19/SARS-CoV-2 data. For the latter, we used the normal and aging lung transcriptome (both bulk RNA-Seq and single cell RNA-seq), protein-protein interactions, and genome-wide association study (GWAS) data.

## MATERIALS and METHODS

### SARS-CoV-2 infection models - Transcriptomic data

We use transcriptomic data from human (Calu-3) and non-human primate (VeroE6) cell lines and from a mouse model (Ad5-hACE2) of SARS-CoV-2 infection. The SARS-CoV-2 infection triggered transcriptome in Calu-3 cell lines is from a recently published study (GSE147507; [15]) wherein 3 samples each were either mock treated or infected with SARS-CoV-2. The second transcriptome signature is based on mRNA profiles of control (mock-infected) and 24h post-SARS-CoV-2-infection (USA-WA1/2020, MOI = 0.3) in Vero E6 cells (kidney epithelial cells extracted from an African green monkey (GSE153940; [16]). The third data set is from a mouse model using Ad5-hACE2-sensitized mice (GSE150847; [17]) that develop pneumonia after infection with SARS-CoV-2, overcoming the natural resistance of mice to the infection.

### Transcriptomic data processing - Differentially expressed genes

Raw data from GSE147507 [15], GSE153940 [16] and GSE150847 [17] were obtained and analyzed using the Computational Suite for Bioinformatics and Biology (CSBB v3.0) [18]. The raw data was downloaded from NCBI Sequence Read Archive (*ProcessPublicData* module) and the technical replicates were merged for individual samples before processing them (*Process-RNASeq_SingleEnd* module). Quality checks [19] and quality trimming [20] were conducted prior to the transcript mapping/quantification step using the RSEM package [21]. Raw counts and transcript per million (TPM) were generated for all samples for further downstream analysis. Within each sample series, differential expression (DE) analysis was carried out based on treatment vs. mock samples using CSBB-Shiny server [22]. RUVSeq [23] was used to remove potential variation and sequencing effects from the data before performing DE analysis using edgeR [24]. Differentially expressed genes (DEGs) were obtained by applying a 1.5 fold change threshold, (i.e., log_2_ *FC* ≥ 0.6 or log_2_ *FC* ≤ −0.6) and a p-value (FDR correction) of <0.05 (Supplementary File 1). For obtaining the human ortholog genes for mouse (Mus musculus) and the green monkey (*Chlorocebus sabaeus*), we used ortholog mappings from the NCBI’s HomoloGene and ENSEMBL databases.

### SARS-CoV-2-Human virus-host protein-protein interactions data

The SARS-CoV-2-human virus-host protein-protein interaction data included a set of 332 human proteins involved in assembly and trafficking of RNA viruses and shown recently through affinity-purification and by mass spectrometry to interact physically with 26 of 29 SARS-CoV-2 structural proteins [25]. These are in addition to the SARS-CoV-2 entry receptor ACE2, and SARS-CoV-2 entry associated proteases, namely, *TMPRSS2*, *CTSB*, and *CTSL*.

### Differentially expressed genes from SARS-CoV-2 infection models and SARS-CoV-2-Human virus-host protein-protein interactome - Network analysis

For all conserved DEGs (genes differentially expressed unambiguously in at least 2 of the 3 transcriptomic sets compared, i.e., Calu-3, VeroE6, or mouse model - Ad5-hACE2) of SARS-CoV-2 infection models and 336 SARS-CoV-2-human PPI genes, we extracted interactions with a score of ≥ 0.9 or experimental interaction score of 0.7 or more from STRING v11 [26]. This DEG-PPI network was clustered (through Cytoscape v3.8.0 [27]) using Markov Clustering Algorithm (MCL) (available as part of the ClusterMaker Plugin [28] in Cytoscape) to identify gene clusters. Briefly, MCL clusters a network to determine modules or clusters of genes with more interactions within the module than to those in the rest of the network [29]. We used the default 2.5 as the inflation factor which determines the granularity (or ‘tightness’) of the clusters and thereby the cluster size. In MCL, each gene can only be assigned to a single module. Modules with 5 or more genes were considered for enrichment analyses while the remaining clusters (with 4 or lesser number of genes) were excluded from further analysis.

### Functional enrichment analysis

Functional enrichment analysis was carried out on the various DEG sets and gene clusters (from MCL clustering) using the ToppFun application of the ToppGene suite [30] and Enrichr [31]. Gene Ontology biological processes, mouse phenotypes, pathways, and 4872 immunologic [32] and 50 hallmark [33] gene sets from MSigDB [34] that are significantly enriched (FDR Benjamini and Hochberg ≤ 0.05) within each of the signature gene sets were clustered together based on their size of enrichments. Similarly, significantly enriched terms from the gene clusters (described above) were integrated and analyzed further. We also performed pathway enrichment analysis using the Elsevier Pathway Collection available as part of the Enrichr application [31].

### Normal adult and aging lung transcriptomics and markers data

To detect specific cell-types potentially perturbed or affected in COVID-19, we intersected the DEGs and gene clusters from SARS-CoV-2 infection models with cell type markers from normal and aging lung. To do this, we compiled cell type marker genes from normal adult human [35–37] and aging mouse lung single cell [38] transcriptomic studies and also bulk RNA-seq data from Genotype-Tissue Expression (GTEx) [39, 40]. We used significant markers (FDR p-value ≤ 0.05; logFC ≥ 0.5) for the enrichment analysis. In case of the aging mouse lung signatures, upregulated (logFC ≤ 0.5) and downregulated (logFC ≥ −0.5) gene sets were used separately for the enrichment analyses. We also generated aging human lung and liver DEG sets from the GTEx data using the BioJupies tool [41]. To do this, we compared samples in the age group 50-79 with those from 20-29 years (lung) and 50-69 years vs. 20-29 years (liver; there were no significant DEGs in 70-79 age group for liver samples). A total of 1683 genes in aging lung (934 upregulated and 749 downregulated) and 745 genes in liver (289 upregulated and 456 downregulated) were significantly dysregulated and were used for the enrichment analysis. Additionally, we also used a curated set of 307 human aging genes from GenAge database [42] (Build February 2020) for enrichment analysis (Supplementary File 2).

### Genome-wide associations (GWA) from PheGenI and GWA studies catalog

To assess the value of our conserved DEGs from SARS-CoV-2 infection models, we intersected them with loci found in GWA studies. To do this we obtained published gene and phenotype trait associations data from the NCBI’s Phenotype-Genotype Integrator (PheGenI) [43] and the NHGRI-EBI GWAS catalog [44]. We used both vulnerability loci of various human physiological traits and the human disease loci. We used an association p-value threshold of 1.0 x 10^−5^ (for PheGenI associations) and excluded all the variants in intergenic regions from the enrichment analysis. We also included child trait associations for the mapped traits from GWAS Catalog. The child terms for each trait were obtained by parsing the experimental factor ontology (EFO) hierarchy [45]. We applied Fisher’s exact test to determine the enrichment.

## RESULTS

### Transcriptomic overlap in *in vitro* and *in vivo* models of SARS-CoV-2 infection

To assess the transcriptomic concordance between in vitro and in vivo models of SARS-CoV-2 infection, we computed pairwise overlaps of the differentially expressed genes (DEGs) from the two in vitro models and the one in vivo model (Table 1). A strong concordance was seen among the upregulated genes and downregulated gene signatures from the three models based on the extent and the significance of the DEG overlaps (Figures 1A, 1B, and 1C). A total of 732 DEGs (537 upregulated and 195 downregulated) were shared between the SARS-CoV-2 infected human Calu-3 and non-human primate VeroE6 cell lines (Figure 1B). Similarly, we found 325 upregulated and 369 downregulated genes common between DEGs from Calu-3 model and Ad5-hACE2-sensitized mice. While there was an overall concordance between the three SARS-CoV-2 infection models, each of the models also had many DEGs unique to the model (Figure 1C). These results highlight the complexity of SARS-CoV-2–host interactions, potential heterogeneity in host response and a dynamic environment driven by the virus replication and the infected cell type.

**Table 1:** List of differentially expressed genes (1.5 log FC; p-value 0.05 FDR) from the 3 SARS-CoV-2 infection models.

| Differentially expressed gene list name | number of DEGs | GSE ID | Reference |
| --- | --- | --- | --- |
| Calu3 SARS-CoV-2 - Downregulated | 2272 | GSE147507 | [15] |
| Calu3 SARS-CoV-2 - Upregulated | 2509 |  |  |
| Mouse Ad5-hACE2 sensitized Mouse - SARS-CoV-2 Downregulated | 2109 | GSE150847 | [17] |
| Mouse Ad5-hACE2 sensitized Mouse - SARS-CoV-2 Upregulated | 1217 |  |  |
| Vero E6 SARS-CoV-2 Downregulated | 953 | GSE153940 | [16] |
| Vero E6 SARS-CoV-2 Upregulated | 1369 |  |  |
| Overall number of unique DEGs: 8286 |  |  |  |

**Figure 1:**
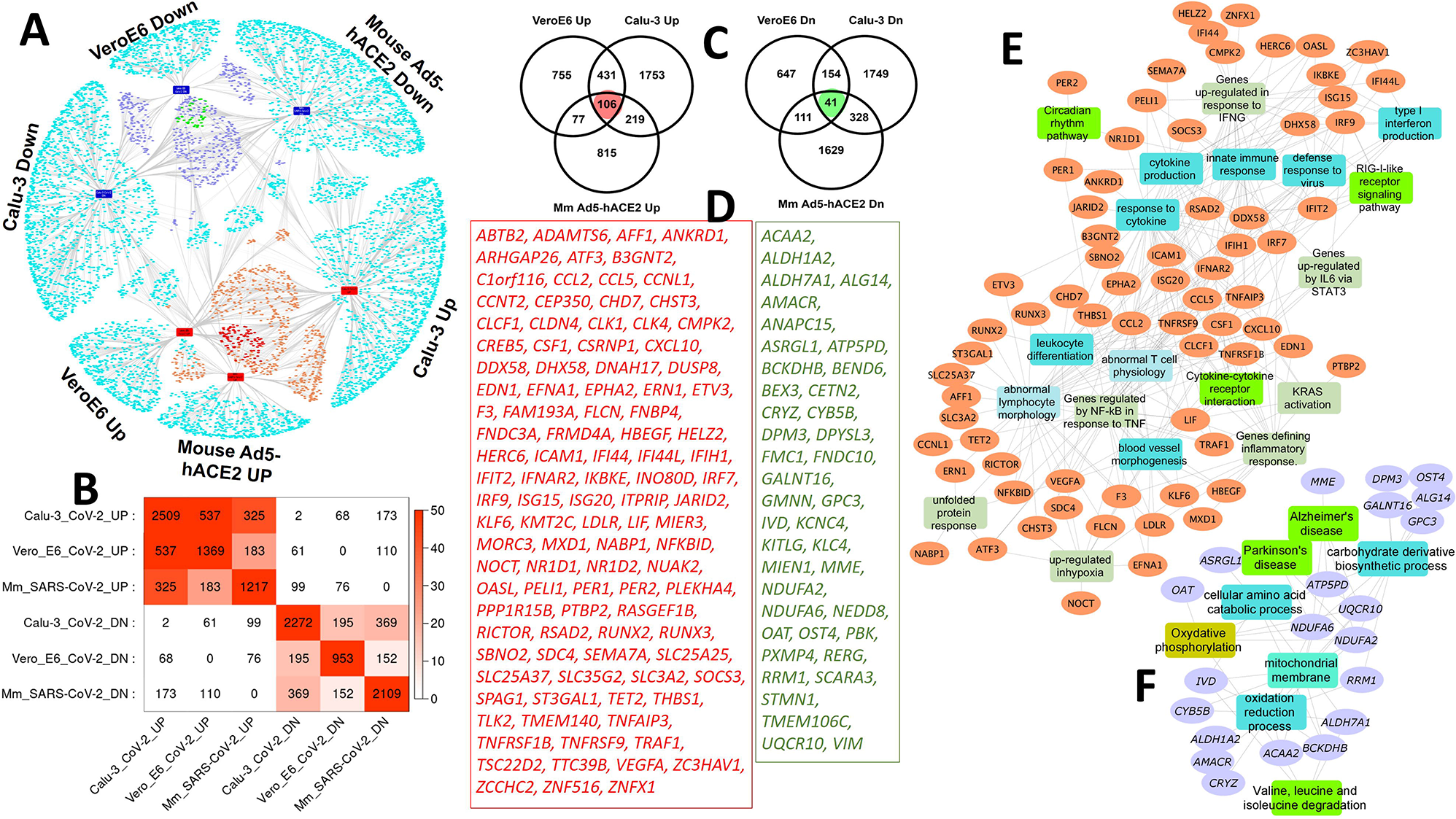
Transcriptomic overlaps among the two in vitro and one in vivo SARS-CoV-2 infection models. **(A)** Network of DEGs from the three SARS-CoV-2 infection models (Calu-3, VeroE6, and Ad5-hACE2 mice). The orange and navy-blue color nodes are genes upregulated or downregulated respectively in at least two models. The red and green color nodes are genes that up- or down-regulated in all the models compared. **(B).** Heatmap indicating the transcriptomic overlaps between the different SARS-CoV-2 infection models. The size and significance of the overlaps is measured using the gene counts and Fisher’s exact test, respectively. **(C)** Venn diagram showing the comparison of the up- or down-regulated differentially expressed genes in the 3 models. **(D)** List of ‘core’ upregulated (106 genes – red font) and downregulated (41 genes – green font) in all the three model systems. **(E) and (F)** Network representation of enriched biological processes and pathways in the core upregulated (panel E) or downregulated (panel F) genes from the SARS-CoV-2 infection models. Orange and purple colored nodes are genes up- or downregulated, respectively. The different colored rectangles are enriched biological processes and pathways. Enrichment analysis was done using the ToppFun application of the ToppGene Suite and network was generated using the Cytoscape application.

### “Conserved” and “Core” transcriptomic signature from SARS-CoV-2 infection in vitro and in vivo models

To obtain a “conserved” transcriptomic signature from the SARS-CoV-2 infection models, we filtered out all ambiguous differentially regulated genes in different models. Specifically, we considered differentially expressed genes in at least 2 of the 3 models compared (i.e., 2 cell lines, namely, transformed lung-derived Calu-3 cells, VeroE6 cells, and mouse model) as conserved SARS-CoV-2 signatures. A total of 1467 conserved genes (833 upregulated and 634 downregulated) were found (Supplementary File 3). There were 106 upregulated and 41 downregulated genes in all the three model systems (Figures 1C and ID) representing the “core” dysregulated transcriptome in SARS-CoV-2 infection. In support of this, these 106 upregulated genes were as expected enriched for innate immune response, cytokine signaling, viral response, RIG-I-like receptor signaling pathway, and leukocyte differentiation (Figure 1E). Additionally, they also showed enrichment for blood vessel morphogenesis, circadian rhythm. The 41 downregulated genes on the other hand were mostly mitochondrial genes and were enriched for metabolic pathways (Figure 1F). Interestingly, there was also an enrichment for neurodegenerative disease pathways (Supplementary File 4).

Functional enrichment analysis of 833 conserved upregulated genes showed enrichment for biological processes and pathways associated with immune system development (119 genes), innate immune response (112 genes), cytokine production (110 genes), response to cytokine (160 genes), cytokine signaling pathway (111 genes) and interferon signaling (30 genes). We also saw genes involved in cardiovascular system development (104 genes), abnormal inflammatory response (173 genes) and abnormal blood cell morphology/development (146 genes). The upregulated signature also had genes (*BHLHE40, NR1D1, PER1, PER2, PER3*) associated with circadian rhythm in mammals (Supplementary File 5). Further, the conserved upregulated genes showed enrichment for several disease pathways such as atherosclerosis, myocarditis, cancers, ROS in vascular inflammation, COPD, pulmonary hypertension, endothelial cell dysfunction, glomerulonephritis, autoimmune disorders, obesity, etc. (Supplementary File 6). Several of these diseases such as cancer, chronic kidney diseases, COPD, obesity, immunocompromised states, cardiovascular diseases are reported as risk factors for severe illness from SARS-CoV-2 infections (https://www.cdc.gov/coronavirus/2019-nCoV/). Multimorbidity is reported to be as high as 35% in people aged 55–64 years and 55% in those aged 65–74 years [46]. The risk of contracting COVID-19 in patients with chronic obstructive pulmonary disorder (COPD), for instance, is reported to be 4-fold higher compared to patients without COPD [47].

The 634 conserved downregulated genes of the SARS-CoV-2 infection models were enriched for several cell cycle associated and other metabolic processes (Supplementary File 5). We also saw enrichment of mitochondrion organization (50 genes) and transport (25 genes). There were more than 60 genes associated with various mitochondrial functions downregulated. Viruses are known to tweak host mitochondrial metabolism to sustain an appropriate viral replication niche [48]. Additionally, several studies have also established a crosstalk between innate immune receptor mediated signaling and the mitochondria [48]. How can SARS-CoV-2 manipulate host mitochondrial morphology and perturb mitochondrial metabolism and how these mechanisms are used to evade the host defense response are some of the outstanding questions that call for further investigation. Equally important, can the immune responses against SARS-CoV-2 be elicited via mitochondria-targeting small intervening agents? Additionally, we found strong enrichments of genes from pathways involved in neurodegenerative diseases such as Alzheimer’s (30 genes), Parkinson’s (27 genes) and Huntington’s (32 genes).

### Normal and aging lung cell type marker enrichment – Conserved transcriptome of SARS-CoV-2 infection models

We next evaluated the 1467 conserved SARS-CoV-2 genes for lung single cell associations by performing enrichment analysis of the 634 downregulated and 833 upregulated DEGs against single cell marker gene sets compiled from three different human lung scRNA-seq studies. Additionally, to explore the correlation between age and COVID-19 morbidity and mortality, we also used aging lung cell markers in mice. Both the upregulated and downregulated gene sets showed enrichment for several epithelial, immune, and stromal cell marker genes (Figure 2A; Supplementary File 7). The marker enrichment also showed concordance between the three models of SARS-CoV-2 infection (Figure 2B). There was a higher incidence of endothelial cell types and its marker genes in the conserved upregulated gene set while the downregulated genes showed enrichment for epithelial cells (proliferating basal, ionocytes, goblet cells) (Figure 2C-G).

**Figure 2:**
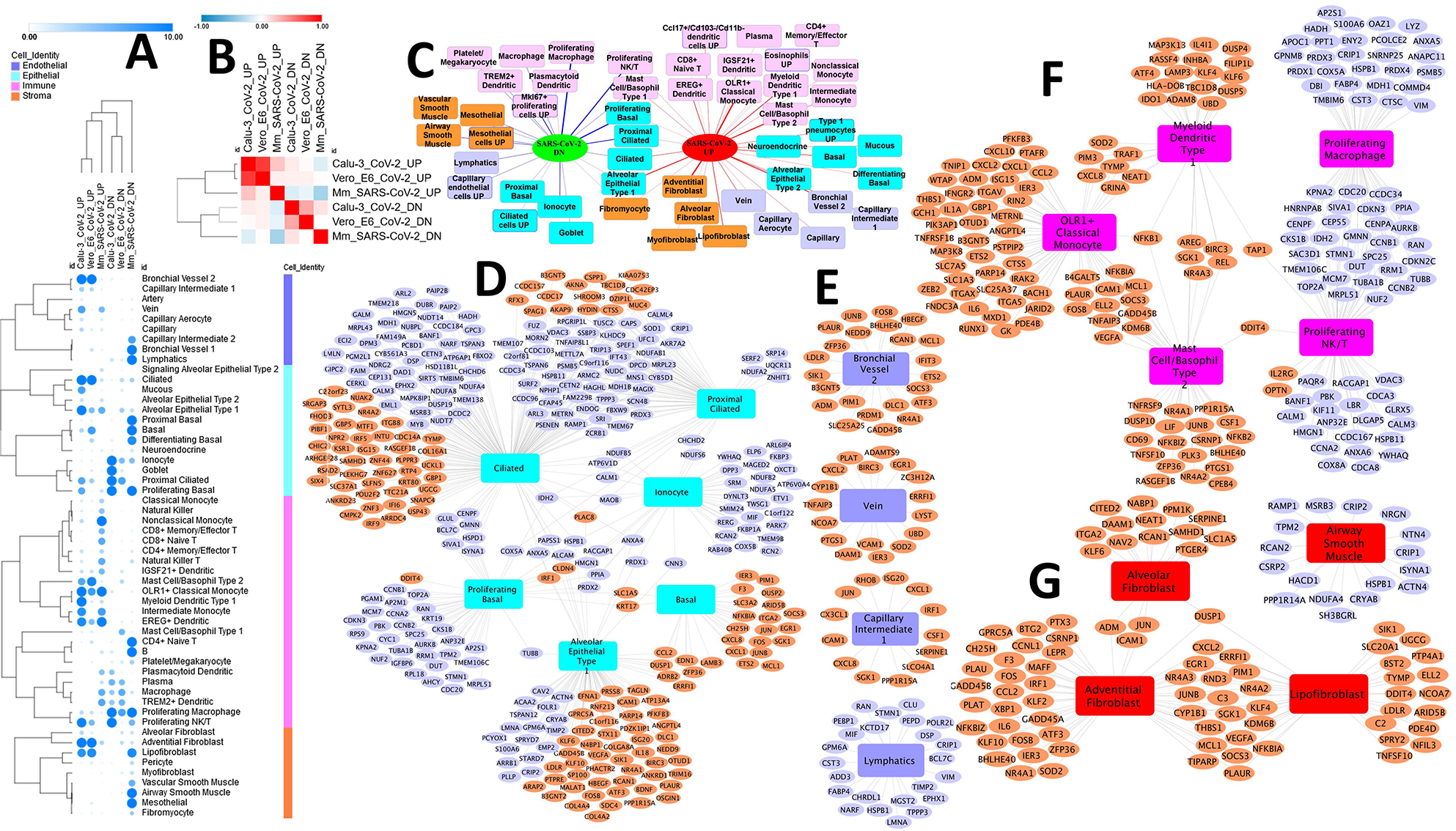
Lung cell type marker enrichment in conserved differentially expressed genes from the 3 SARS-CoV-2 infection models. **(A)** Enrichment heatmap of single cell markers from normal human lung [35] among the differentially expressed genes from the 3 models of SARS-CoV-2 infection. The size and intensity of the colors in the circles is proportional to the significance of enrichment as measured by Fisher’s exact test (negative log p-values). **(B)** Heatmap showing the correlation between the three SARS-CoV-2 infection models based on the shared enriched cell markers. **(C).** Network of enriched cell types from normal human lung [35] and aging mice [38] in the conserved differentially expressed genes from the 3 models of SARS-CoV-2 infection. The different colored rectangles are various cell types. The pink colored rectangles are immune cell types, purple colored ones are endothelial, blue colored rectangles are epithelial, and the orange colored nodes are stromal cell types. The width of the edges is proportional to the significance of the cell type enrichment (negative log p-value) measured by Fisher’s exact test. **(D)-(G)** Network representation of enriched cell types and associated conserved differentially expressed genes from the 3 SARS-CoV-2 infection models. Conserved upregulated genes are in orange while the downregulated genes are in purple. Panels D, E, F, and G represent epithelial, immune, endothelial, and stromal cell type enrichment networks, respectively.

Conserved upregulated genes were primarily enriched for both classical (64 genes) and nonclassical (28 genes) monocyte markers (Figure 2F; Table 2). They also showed exclusive enrichments for endothelial cells in addition to a significant number of adventitial (52 genes) and lipofibroblast (37 genes) markers from [35] (Figure 2G). Furthermore, alveolar epithelial type 1 (AT1) gene markers from both normal and aging lung samples were enriched in the upregulated genes (Figures 2D) in addition to several endothelial cell markers (Figure 2E). Interestingly, conserved downregulated genes were enriched for markers from proliferating macrophages (58 genes), NK/T cells (51 genes) and lymphatic cells (24 genes) in normal lung (Figure 2F). They also had many epithelial markers, associated with cilia (121 genes), proximal cilia (64 genes) and ionocytes (39 genes). Interestingly, the conserved downregulated genes were enriched for proliferating basal cell markers (46 genes) while the upregulated signature had genes (28 genes) expressed in normal basal cells (Figure 2D). A recent study reported that distinct cell populations proliferate in different regions of the respiratory system following SARS-CoV-2 infection [49].

**Table 2:**
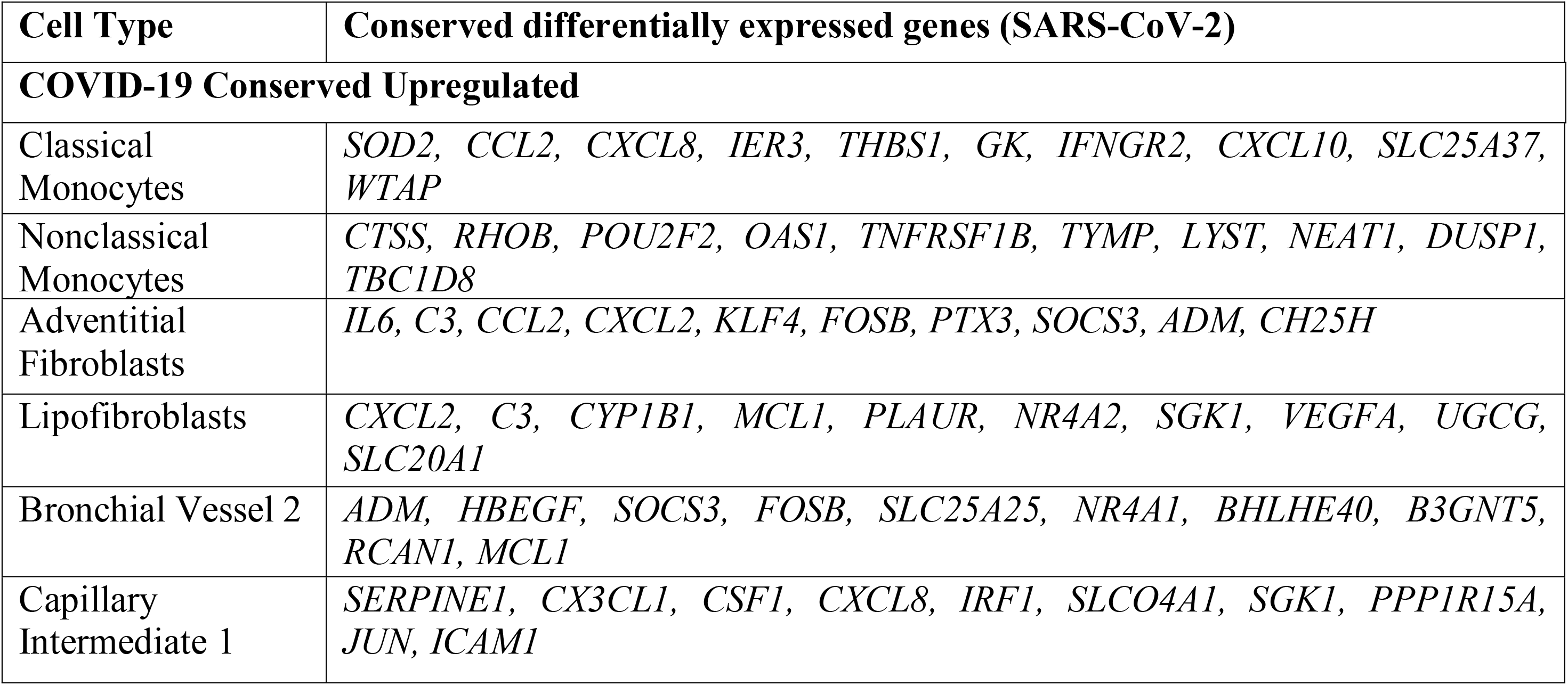
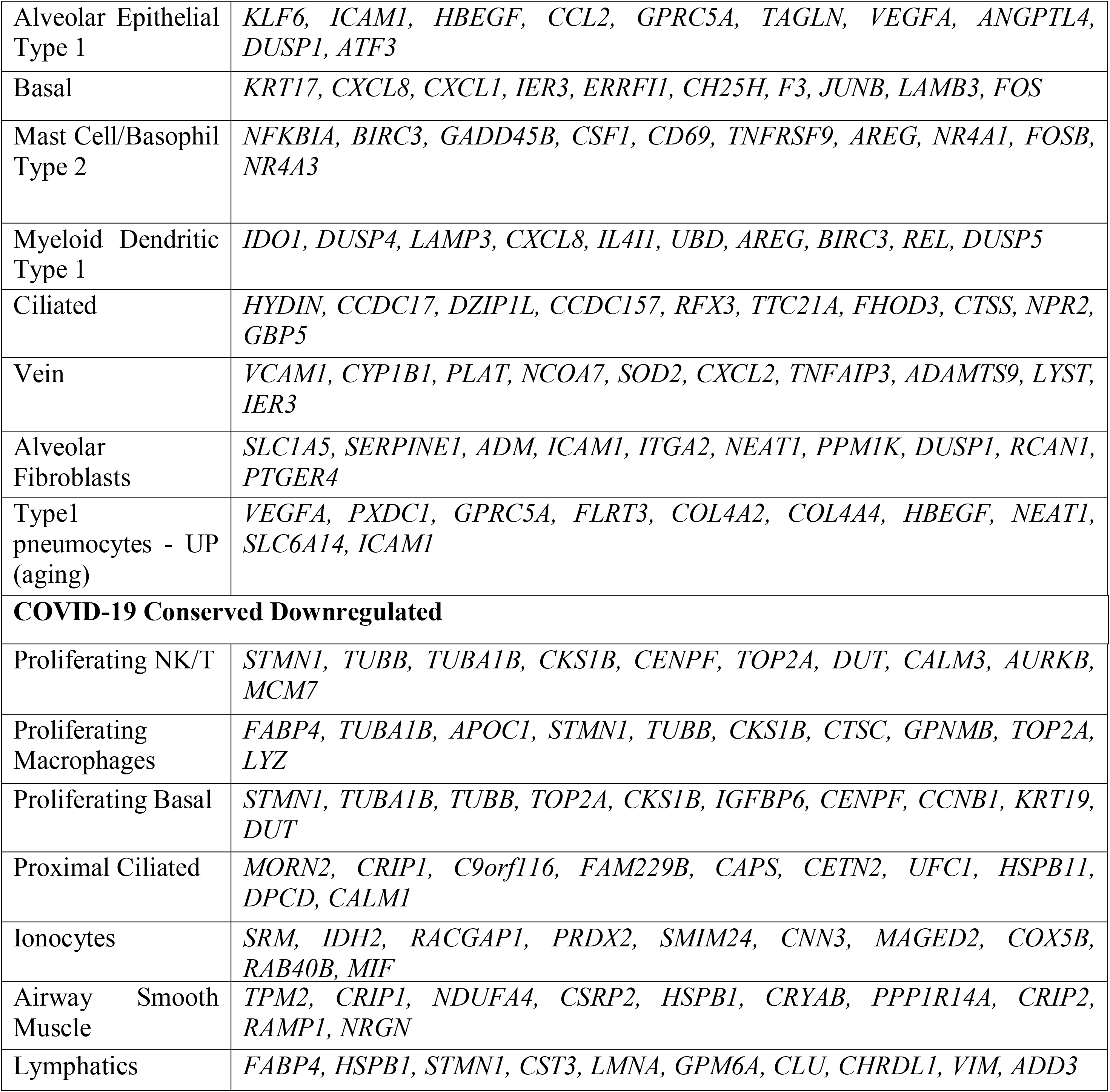
List of enriched normal and aging lung cell types and their marker genes which are differentially expressed in conserved DEGs of SARS-CoV-2 infection models (top 10 most significantly associated marker genes are listed). Complete set of enrichment results can be found in Supplementary File 7.

Similar cell marker enrichment trends were seen in the core SARS-CoV-2 signature (147 genes dysregulated in all the three model systems). The upregulated gene set included markers from classical monocyte (14 genes) and alveolar epithelial type 1 (15 genes) cells while the downregulated genes were enriched for proliferating basal and NK/T cell type markers.

The conserved gene signature also displayed significant overlap with DEG sets from aging liver and lung. SARS-CoV-2 conserved upregulated genes were enriched for genes overexpressed (56 genes) in aging lung tissue while the downregulated gene set was enriched (57 genes) for genes that are downregulated in aging lung. This overall positive concordance between COVID-19 and aging lung agrees with the fact that aging is one of the risk factors for COVID-19 morbidity and mortality. Similarly, there were 43 genes (*ARHGAP31, CDC42EP2, CCL2, NCEH1, RHCG, ABCC1, SLC1A5, CD274, AGO2, NOS3, NPC1, GPRC5A, HAVCR2, GPR176, PDE4B, ARHGAP30, KLF4, KRT80, PLAU, PLAUR, IL4I1, THEMIS2, CREB5, IL6, IL11, METRNL, C15orf39, ITGA2, PTAFR, ITGA5, ITGAM, ITGAX, PTGS1, PTPRE, PTX3, ZSWIM4, RHBDF2, HBEGF, DUSP4, DUSP5, LGALS9, FOSL1 and LIF*) that are downregulated in aging liver but were found to be upregulated in COVID-19. While these were immune response and apoptotic genes, the presence of *IL11*, *PLAU* and *PLAUR* is interesting. *IL11*, a hematopoietic cytokine and member of *IL6* family of cytokines, is known to have potent thrombopoietic activity. The *PLAU-PLAUR* (uPA-UPAR) system is reported to contribute to fibrinolysis, inflammation, innate and adaptive immune responses, and tissue remodeling [50, 51]. *PLAU-PLAUR* are known to take part in the process of coagulation disorder in patients with systemic inflammatory response syndrome (SIRS) and increased levels of *PLAUR* are reported to promote the development of SIRS to multiple organ dysfunction syndrome (MODS) [52]. Further, retrospective studies [53, 54] have also shown that the measurement of soluble uPAR (suPAR) levels in serum, tissue and urine of patients can predict disease severity and outcome. Intriguingly, *SERPINE1*, the gene encoding PAI-1, the endogenous inhibitor of *PLAU-PLAUR* system was also found to be upregulated in COVID-19. Further studies are needed to understand the molecular mechanisms that promote the expansion of distinct cell subpopulations that may be involved in regenerating a functional respiratory system in COVID-19 patients with severe pneumonia. Finally, the conserved upregulated DEGs showed an enrichment for human aging genes from GenAge (benchmark database of aging genes – Build February 2020) with an overlap of 32/833 (p-value <0.05) genes.

### Conserved DEGs and genetic risk factors - PheGenI/GWAS Catalog trait enrichment

The vast majority of SARS-CoV-2 infected individuals experience mild or no symptoms and mortality in a subset of patients is primarily due to acute respiratory distress syndrome, bilateral interstitial pneumonia followed by severe respiratory failure. Earlier studies have reported association of mortality with older age, especially males, and comorbid conditions including obesity, cardiovascular diseases, hypertension, and diabetes. The presence of other genetic risk factors cannot be however ruled out given that we are still amid the pandemic. Furthermore, finding the genetic risk factors can potentially enable not only in the identification of vulnerable people but also in deeper understanding of the mechanisms underlying the outcomes or sequelae of the disease. A recent study discovered 5 risk variants (4 genes – *ALOXE3, ERAP2, BRF2*, and *TMEM181*) as potentially significant risk factors for death in COVID-19 patients [55]. The gene *ALOXE3* was among the 833 conserved upregulated genes. To assess the impact of other traits more generally, both physiological and pathological, we performed an enrichment analysis of the conserved DEGs of SARS-CoV-2 infection models against all human genotype-phenotype associations from the Phenotype-Genotype Integrator (PheGenI) [43] and GWAS Catalog [44] databases (Supplementary Files 8 and 9).

Among the top enriched traits, were platelet functions, blood pressure, body mass index, respiratory function tests, cholesterol and triglycerides, and several neurodegenerative and neuropsychiatric diseases (Table 3). The associated genes were either up- or down-regulated in SARS-CoV-2 infection models (Figures 3A-J). We also found enrichment for smoking and chronic obstructive pulmonary disease. The findings from the recent OpenSAFELY (https://opensafely.org/) study [56], based on ~17 million COVID-19 patients’ detailed primary care records lend support to most of our results. For instance, among the top enriched traits were blood pressure (70 genes; 34 upregulated and 36 downregulated) and body mass index (73 genes; 37 upregulated and 36 downregulated), in addition to platelet function tests (157 genes; 112 upregulated and 45 downregulated) and lung function measurements (52 genes; 24 upregulated and 28 downregulated). The other top enriched traits were cholesterol, triglycerides (50 genes; 34 upregulated and 16 downregulated), and smoking (33 genes; 13 upregulated and 20 downregulated) (Table 3). Surprisingly, we also found enrichment for child developmental disorders and attention deficit disorder with hyperactivity. *TRANK1* (tetratricopeptide repeat and ankyrin repeat containing 1), one of the genes associated with child developmental disorder has been shown to be an interferon-stimulated gene (ISG) in both humans and *P. vampyrus* (large fruit bat) cells. We found this gene upregulated in our study. A recent study reported that a polymorphism in the *TRANK1* region affects brain development in the presence of a perinatal injury, with pathophysiological consequences such as KLS (Kleine-Levin Syndrome), bipolar disorder and schizophrenia [57]. KLS, a rare disease affecting adolescents is characterized by relapsing-remitting episodes of severe hypersomnia, cognitive impairment, and behavioral disturbances [58, 59]. Interestingly, although the etiology of KLS is still unknown, a strong positive correlation between upper respiratory infections or flu-like illness and symptomatic episodes of KLS has been found [60, 61].

**Table 3:**
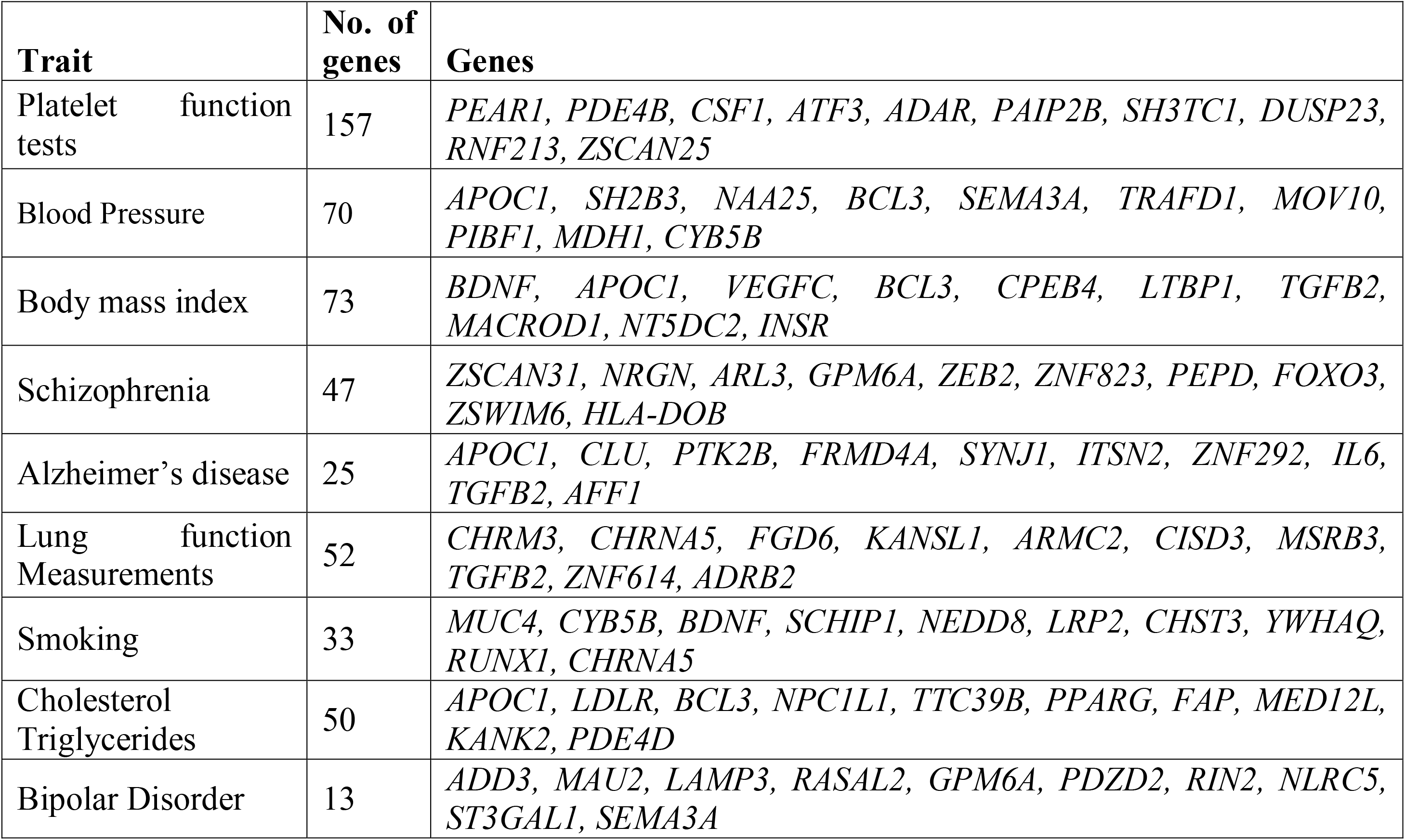
Physiological and pathological GWA trait enrichment in SARS-CoV-2 DEG. Top enriched traits, and the ten most significantly associated conserved DEGs from the 3 SARS-CoV-2 infection models are shown here (see Supplementary Files 8 and 9 for the complete list and more details)

**Figure 3:**
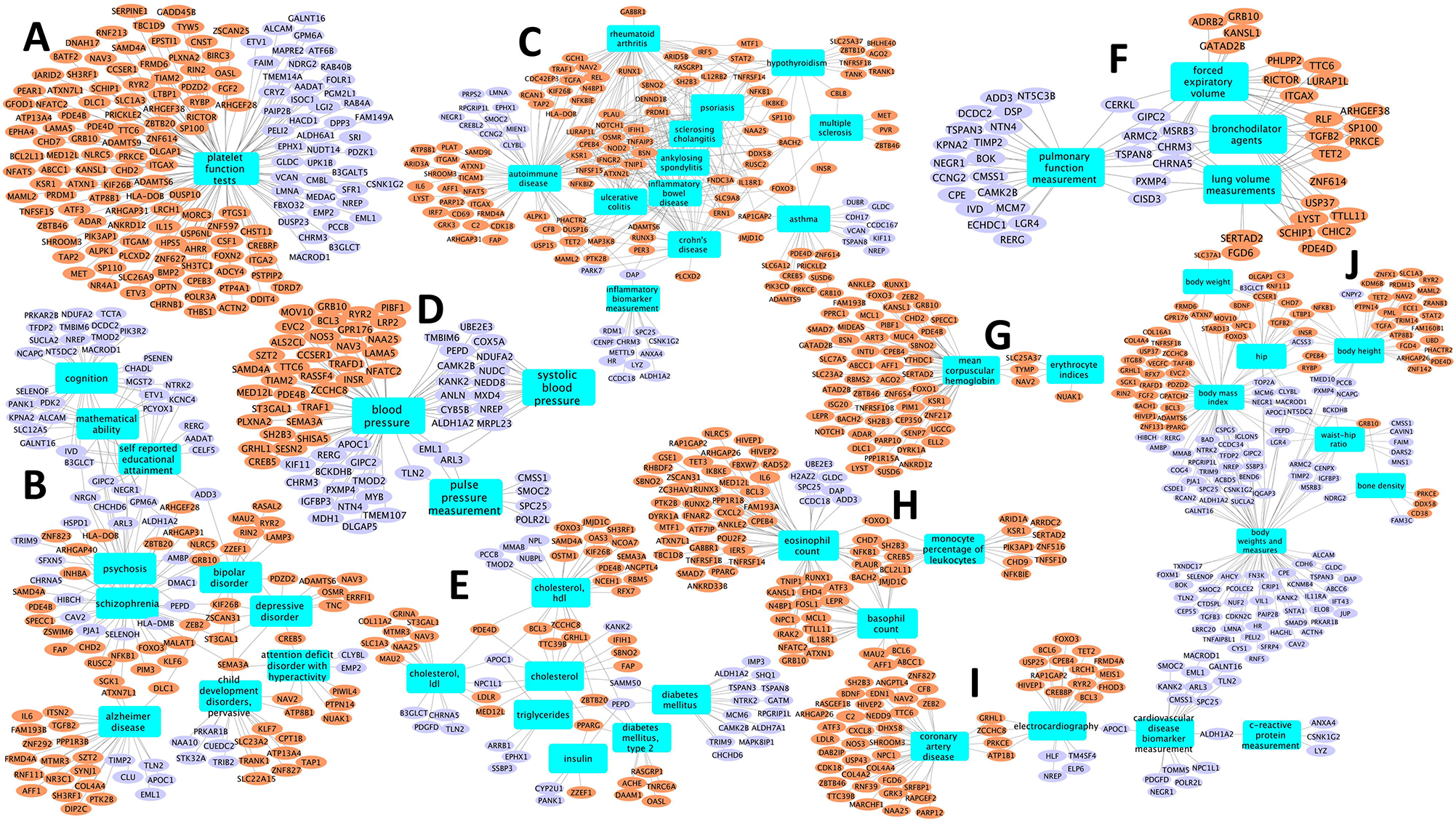
GWA loci enrichment in conserved differentially expressed genes from the 3 SARS-CoV-2 infection models. Network representation of select enriched PheGenI/GWA traits and their associated conserved DEGs from the 3 models of SARS-CoV-2 infection. Upregulated genes are in orange, downregulated genes are in purple, and the enriched traits are shown as blue colored rectangles. Networks are shown as different panels (A-J) based on their broad categorization, where possible. For instance, panel B shows cognition, neurodegenerative and neuropsychiatric GWA loci with their associate genes while panels C and F represent immune disorders and lung function, respectively.

On similar lines, we found enrichment for other cognition impairment and neurodegenerative disorders: Alzheimer’s (25 genes; 20 upregulated and 5 downregulated), Schizophrenia (47 genes; 29 upregulated and 18 downregulated), and Bipolar (13 genes; 11 upregulated and 2 downregulated). Both acute and delayed neurological and neuropsychiatric effects have been associated with previous viral pandemics [62]. For instance, earlier studies reported infection with SARS-CoV-1 as a risk factor for developing Parkinson’s disease [63]. Further, results from autopsy of COVID-19 patients showed edema and hyperemia in brain along with neuron degeneration [64]. However, long-term neurological and neuropsychiatric sequelae for SARS-CoV-2 infection are currently unknown and may only be revealed in coming months or even years. Even though the role of immune dysregulation in classical autoimmune brain diseases (e.g., multiple sclerosis, autoimmune encephalitis), psychiatric disorders (e.g., schizophrenia, autism spectrum disorder, bipolar disorder, and depression) is well-documented [65] there are relatively few reports of viral infections as risk factors for psychiatric disorders [66, 67]. We also found a strong enrichment for sleep and sleep measurement traits (19 genes) among the downregulated genes. In case of COVID-19, the unique psychosocial stressors associated with social distancing may further add to the neuropsychiatric symptom burden. Although, a recent study assessing the neuropsychiatric presentations of SARS, MERS, and COVID-19 using systematic reviews and meta-analysis found little evidence to support the plausibility of neuropsychiatric complications as COVID-19 sequelae, given the early stages of this pandemic, it is too early to rule out the long-term neuropsychiatric effects. Unlike SARS-CoV, SARS-CoV-2 was reported to modestly replicate in neuronal (U251) cells [68], lending further credence to the possibility that this virus can lead to neurological manifestations in the COVID-19 patients. Furthermore, beta coronavirus (HCoV-OC43) has been previously associated with fatal encephalitis in an 11-month-old boy with severe combined immunodeficiency (SCID) [69]. Prospective monitoring of COVID-19 patients including children and younger adults who have recovered from COVID-19 must determine potentially long-term neuropsychiatric outcomes. Further exploration of our findings with regards to their utility in mechanistic understanding and characterizing of COVID-19, and their potential in clinical risk profiling of COVID-19 patients is warranted.

### Differential expression of SARS-CoV-2 host interacting genes during infection

To see if any of the genes encoding SARS-CoV-2 host interacting proteins [25] (336 genes) are differentially expressed during COVID-19 infection, we compared SARS-CoV-2-human interaction map with the SARS-CoV-2 conserved signature (833 upregulated and 634 downregulated). About 51% (171/336) of the SARS-CoV-2-human interacting proteins were differentially expressed in at least one of the 3 model systems (Figure 4). Of these, 29 genes (16 upregulated and 13 downregulated) were part of the conserved signature while 3 genes (two upregulated - *PTBP2* and *CEP350* and one downregulated - *CYPB5B*) were part of the core signature (Figure 4A; Table 4).

**Table 4:** List of 29 differentially expressed genes in SARS-CoV-2 infection models that encode proteins interacting with SARS-CoV-2 proteins. 16 upregulated and 13 downregulated genes in SARS-CoV-2-triggered transcriptome whose encoded proteins interact directly with SARS-CoV-2 proteins are presented along with their function and potential druggability [25]. Host genes highlighted in bold (and with asterisk) are potential drug candidates.

| **Host Gene** | **SARS-CoV-2 gene** | **Biological function of COVID-19 Gene** |
| --- | --- | --- |
| *MOV10* | N | Signaling |
| ***ABCC1*\*** | Orf9c | Mitochondria |
| *ZC3H7A* | Nsp12 |  |
| *ATP1B1* | M | Vesicle trafficking |
| *MYCBP2* | Nsp12 |  |
| *PTBP2* | Orf9b | Signaling |
| *ZNF503* | Nsp9 | Extracellular matrix |
| *TMPRSS2* | Spike protein | Transmembrane protease |
| *PLAT* | Orf8 | Vesicle trafficking |
| *CSNK2A2* | N | Signaling |
| *CEP350* | Nsp13 | Epigenetic and gene expression regulators |
| *PVR* | Orf8 | Vesicle trafficking |
| *INHBE* | Orf8 | Vesicle trafficking |
| *CLIP4* | Nsp13 | Epigenetic and gene expression regulators |
| *SNIP1* | N | Signaling |
| *AKAP9* | Nsp13 | Epigenetic and gene expression regulators |
| *CENPF* | Nsp13 | Epigenetic and gene expression regulators |
| *AP2M1* | Nsp10 | Vesicle trafficking |
| *NPC2* | Orf8 | Vesicle trafficking |
| *PPT1* | Orf10 | Ubiquitin ligases |
| *PRKAR2B* | Nsp13 | Epigenetic and gene expression regulators |
| *CISD3* | Orf8 | Vesicle trafficking |
| *ATP6AP1* * | Nsp6 | Vesicle trafficking |
| *CLCC1* | Orf3a | Vesicle trafficking |
| *CSDE1* | Orf9b | Signaling |
| *ELOB* * | Orf10 | Ubiquitin ligases |
| *SIRT5* | Nsp14 |  |
| *CYB5B* | Nsp7 | Vesicle trafficking |
| *TIMM8B* | Orf10 | Ubiquitin ligases |

**Figure 4:**
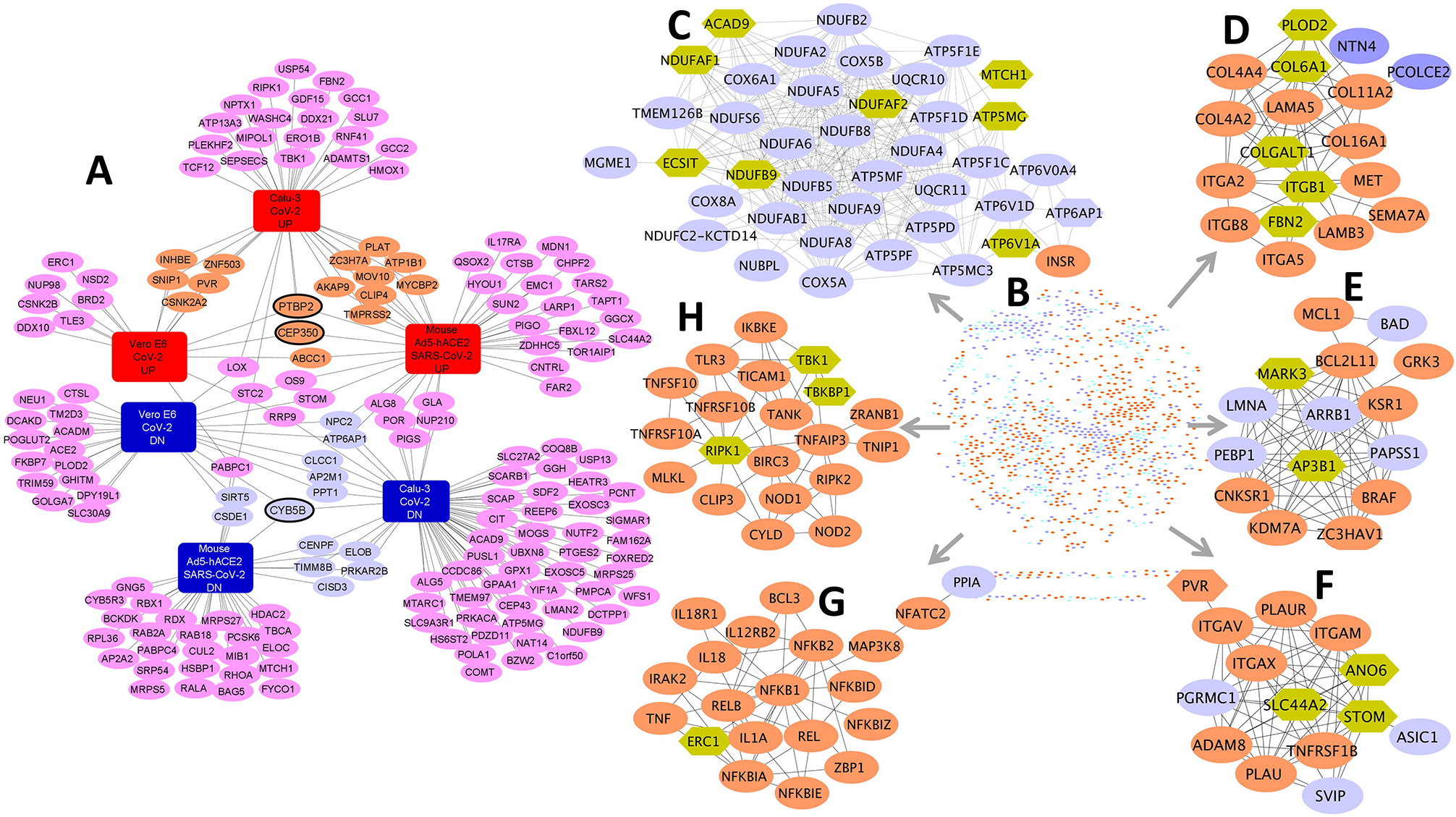
Network of conserved differentially expressed genes of SARS-CoV-2 infection models and SARS-CoV-2-host protein-protein interactions. **(A)** Network of 176 differentially expressed genes of three SARS-CoV-2 infection models that encode proteins interacting with SARS-CoV-2 viral proteins. The orange and purple colored nodes are conserved up or downregulated genes respectively while the 3 genes with blue border (PTBP2, CEP350, and CYB5B) are part of the core set of genes. **(B)** STRING-based interaction network of conserved DEGs and SARS-CoV-2-human viral-host protein interactome. **(C)-(F).** Example gene clusters from the conserved DEG and SARS-CoV-2-host integrated interactome. Clusters (based on MCL network clustering) shown are C-5 (Panel C), C-9 (Panel D), C-11 (Panel E), C-13 (Panel F), C-7 (Panel G) and C-8 (Panel H). Conserved up and downregulated genes are in orange and purple colors respectively while the hexagons are part of the SARS-CoV-2-host protein interactome that are not differentially expressed but directly interact with the conserved DEGs from SARS-CoV-2 infection models.

### SARS-CoV-2 infection model DEGs – Interacting gene clusters

It is well proven that viral infection is tightly associated with host protein complexes which are manipulated by viruses to hijack the host cell biological processes. To better understand this phenomenon in COVID-19, we next combined the conserved transcriptomic signature (833 upregulated and 634 downregulated genes) from the three SARS-CoV-2 infection models with SARS-CoV-2-human interaction map [25] (336 genes). We then queried the STRING (v11) [26] database using this combined set of genes to build a SARS-CoV-2 DEG and interaction map. STRING database collects the available PPI information from multiple sources and assigns a confidence score by considering both direct and indirect interactions. Using highest confidence score (0.9) or experimental interaction score of 0.7 or more, we aimed to filter out the PPIs with lower probability to be interactions. With this criterion, there was an enrichment for PPIs (p-value < 1.0e-16) among the SARS-CoV-2 DEGs and SARS-CoV-2-interacting proteins. In other words, the combined set of SARS-CoV-2 conserved signature and SARS-CoV-2-human interaction map (1080 nodes; 5481 edges; Figure 4B) have significantly more interactions among themselves than what would be expected for a random set of proteins of similar size, drawn from the genome. Since two or more interacting proteins have a higher probability of sharing similar function, we next extracted network clusters from this joint interactome using Markov clustering (MCL) algorithm, with the default inflation parameter (2.5). In total, we found 153 clusters of varying gene counts (Supplementary File 10). Biological significance and COVID-19 relevancy of these clusters was inferred through enrichment analysis of the clusters for normal and aged lung cell markers and enrichment for the SARS-CoV-2-human interaction map proteins. We selected 35 candidate clusters with each having at least 5 genes. These 35 clusters were made up of a total of 797 genes of which 627 were conserved DEGs in SARS-CoV-2 infection models. Furthermore, 186/797 genes encoded proteins that directly interact with SARS-CoV-2 proteome, and 16/797 genes were conserved DEGs and also part of SARS-CoV-2-host protein-protein interactions (See Figure 4C-H for six example clusters and Supplementary File 10 for more details). Of the 35 clusters, 29 clusters had at least one gene encoding protein that interacts with SARS-CoV-2 proteome. We hypothesize that these SARS-CoV-2 targeted human protein clusters could not only be informative in deciphering the COVID-19 pathophysiology but also be useful to infer the function of the SARS-CoV-2 target based on other members in this SARS-CoV-2 targeted human protein clusters.

### Gene clusters from DEGs of SARS-CoV-2 infection models and SARS-CoV-2-human protein interactome to characterize COVID-19

#### Gene clusters - lung single cell markers

Of the 35 selected gene clusters, 17 clusters were enriched for at least one lung cell type (normal or aging lung) (Figure 5A). These 17 gene clusters made up 633 genes (259 marker genes) of which 500 were conserved DEGs from SARS-CoV-2 infection models and 147 were from SARS-CoV-2-host interacting proteins (Supplementary File 11). We saw strong concordance among the cell markers between different clusters (Figures 5B-D). For example, Cluster C-1 (190 genes) and Cluster C-2 (92 genes) consisted of marker genes for alveolar epithelial type 1, proliferating macrophages, and adventitial fibroblasts (Figure 5A) (Table 5). These clusters also shared strong enrichment for genes that are downregulated in human aging lungs (DEGs from GTEx). Cluster C-9 (18 genes) consisted of gene markers in fibroblasts, myofibroblasts and smooth muscle cells and shared enrichments with clusters C-1, C-2, and C-3 (Figure 5B). Ionocyte cell marker [35] genes were confined to Clusters C-5 (40 genes; 12 markers) and C-22 (8 genes; 3 markers) (Figure 5C) while Cluster C-10 was enriched for endothelial cell types and shared enrichments with C-2 (Supplementary File 11).

**Table 5:** Candidate clusters in SARS-CoV-2 DEG and interaction map along with their enriched normal and aging cell types. Clusters with ≥ 5 genes enriched for at least one normal and/or aging lung cell type are shown here. Complete list of clusters (Supplementary File 10), their enriched cell types (Supplementary File 11) are made available as Supplementary data.

| **Cluster** | **Enriched cell markers** |
| --- | --- |
| C-1 (190 genes) | Proliferating NK/T, Proliferating Basal, Mki67+ proliferating cells-UP (aging), Proliferating Macrophage, Adventitial Fibroblasts, alveolar epithelial type 1 |
| C-2 (91 genes) | Proliferating Epithelial, Ciliated, Ciliated cells-DN (aging), Proliferating Macrophage, Classical Monocytes, Alveolar epithelial type 1, adventitial fibroblasts |
| C-3 (81 genes) | Adventitial Fibroblasts, Lipofibroblasts, Bronchial Vessel 2, Classical Monocytes, Mast cells |
| C-4 (73 genes) | Ciliated, Capillary endothelial cells |
| C-5 (40 genes) | Ionocytes, Proximal Ciliated |
| C-6 (34 genes) | Bronchial Vessel 1, Lipofibroblasts, Mesothelial |
| C-7 (20 genes) | Dendritic cells, Mast cells, Ccl17+/Cd103-/Cd11b-dendritic cells-UP |
|  | (aging) |
| C-9 (18 genes) | Alveolar Epithelial Type 1, Fibroblasts, Basal, Myofibroblasts, Smooth Muscle Cells |
| C-10 (17 genes) | Lymphatic, Peri bronchial, Arterial |
| C-11 (15 genes) | Dendritic, Mast cells |
| C-13 (14 genes) | Classical Monocytes |
| C-18 (10 genes) | Proliferating Epithelial, Proliferating T-Cells |
| C-22 (8 genes) | Ionocytes, Macrophages, Proliferating T-Cells |
| C-30 (6 genes) | Proliferating T-Cells |
| C-31 (6 genes) | Arteries |
| C-34 (5 genes) | Plasma cells |
| C-35 (5 genes) | Plasma cells |

**Table 6:** Enriched functional terms (biological processes, pathways, cell types, and phenotypic traits) shared among the 35 candidate gene clusters from the integrated network of conserved DEGs of the 3 SARS-CoV-2 infection models and the SARS-CoV-2-host protein interaction map. The shared enriched terms are found through meta-analysis of the enriched terms from different annotation categories for the 35 gene clusters. The complete network along with all enriched terms and cluster details are presented in the Supplementary File 15.

| Cluster | Enriched functional terms |
| --- | --- |
| C-1 (190 genes) | Innate immune response; response to type I interferon; Cell cycle; Proliferating NK/T; Proliferating macrophages; AT1; Adventitial fibroblasts; Platelet function tests |
| C-2 (91 genes) | Toll-like receptor signaling pathway; Abnormal blood vessel morphology; Arteriosclerosis; Goblet; AT2; Capillary; C-X-C motif chemokine 11 measurement |
| C-3 (81 genes) | DNA-templated transcription, initiation; RNA splicing; abnormal heart ventricle morphology; Basal; Fibroblasts; Adventitial fibroblasts; monocytes; macrophages |
| C-4 (73 genes) | Golgi vesicle transport; membrane docking; Ciliated; Capillary; abnormal neural tube ventricular layer morphology |
| C-5 (40 genes), C-12 (14 genes) and C-28 (6 genes) | Mitochondrion organization; mitochondrion support; oxidative phosphorylation; ATP biosynthesis; abnormal mitochondrial crista morphology; Alzheimer's; Parkinson's |
| C-6 (34 genes) | RNA catabolic process; RNA degradation; Bronchial Vessel 1; CD4+ Naive T; Proximal Basal |
| C-7 (20 genes) and C-8 (20 genes) | Response to cytokine; cytokine production; T cell receptor signaling pathway; abnormal interleukin secretion; Mast Cell/Basophil Type 2; dendritic cells; ulcerative colitis; Crohn's disease; inflammatory biomarker measurement |
| C-9 (18 genes) | Cell-substrate adhesion; Genes encoding collagen proteins; ECM; Endothelial cells; AT1; Fibroblasts; Myofibroblasts; Smooth muscle cells |
| C-10 (17 genes) | cell-substrate adhesion; platelet degranulation; vesicle fusion; VEGF and VEGFR signaling network; Smooth muscle cells; Lymphatic cells |
| C-11 (15 genes) | Apoptotic mitochondrial changes; PAR1-mediated thrombin signaling events; Ceramide signaling pathway; abnormal melanocyte morphology; abnormal splenocyte apoptosis; leukocyte counts |
| C-13 (14 genes) | neutrophil activation; neutrophil mediated immunity; liver inflammation; OLR1+ Classical Monocytes |
| C-14 (12 genes) | chromosome organization; histone modification; negative regulation of RNA biosynthetic process |
| C-15 (11 genes) | trans-synaptic signaling by neuropeptide; Notch signaling pathway; type II diabetes mellitus; influenza A severity measurement |
| C-16 (11 genes) and C-17 (11 genes) | fatty acid catabolic process; Tryptophan metabolism; Genes up-regulated during adipocyte differentiation; |
| C-18 (10 genes) | DNA replication; Pyrimidine metabolism; Cell cycle; Proliferating NK/T; Proliferating macrophages; Proliferating Basal |
| C-19 (9 genes) | cytoplasmic translation; Aurora A signaling; Canonical NF-kappaB pathway |
| C-20 (9 genes) | glycoprotein metabolic process; aminoglycan metabolic process; heparan sulfate/heparin (HS-GAG) metabolism; |
| C-21 (8 genes) | Pyrimidine metabolism; genes involved in DNA repair; decreased coronary flow rate; abnormal renal vascular resistance; Airway wall thickness measurement; orotic acid measurement |
| C-22 (8 genes) | cellular detoxification; oxidation-reduction process; abnormal mitochondrial physiology; neuron degeneration; Huntington's disease; Ionocyte; Proximal ciliated; Proliferating T cells; |
| C-23 (8 genes) and C-35 (5 genes) | Translation elongation; mitochondrial gene expression; regulation of translation; Plasma cells |
| C-24 (7 genes) | regulation of cellular amine metabolic process; Aurora A signaling; increased mean corpuscular hemoglobin concentration |
| C-25 (7 genes) | circadian rhythm; p38 signaling mediated by MAPKAP kinases; abnormal brain wave pattern |
| C-26 (7 genes) | Cell cycle; cell junction organization; pulmonary edema; hemopericardium; Arrhythmogenic right ventricular cardiomyopathy (ARVC) |
| C-27 (7 genes) | Golgi vesicle transport; endosomal transport; endomembrane system organization |
| C-29 (6 genes) | liver development; hormone-mediated signaling pathway; Adherens junction; Atypical NF-kappaB pathway; autoimmune disease |
| C-30 (6 genes) | cellular response to corticosteroid stimulus; Insulin-mediated glucose transport; mTOR signaling pathway; ProliferatingT Cells |
| C-31 (6 genes) | retinol metabolic process; Genes encoding proteins involved in metabolism of fatty acids; Artery; Cystic fibrosis |
| C-32 (6 genes) | Cellular amino acid metabolic process; glycine metabolic process; decreased urine creatine level; abnormal amino acid level; urinary potassium to creatinine ratio |
| C-33 (5 genes) | activin receptor signaling pathway; RNA degradation; TGF-beta signaling pathway; Calcium signaling in the CD4+ TCR pathway |
| C-34 (5 genes) | ERAD pathway; Hypoxic and oxygen homeostasis regulation of HIF-1-alpha; Plasma cells |

**Figure 5:**
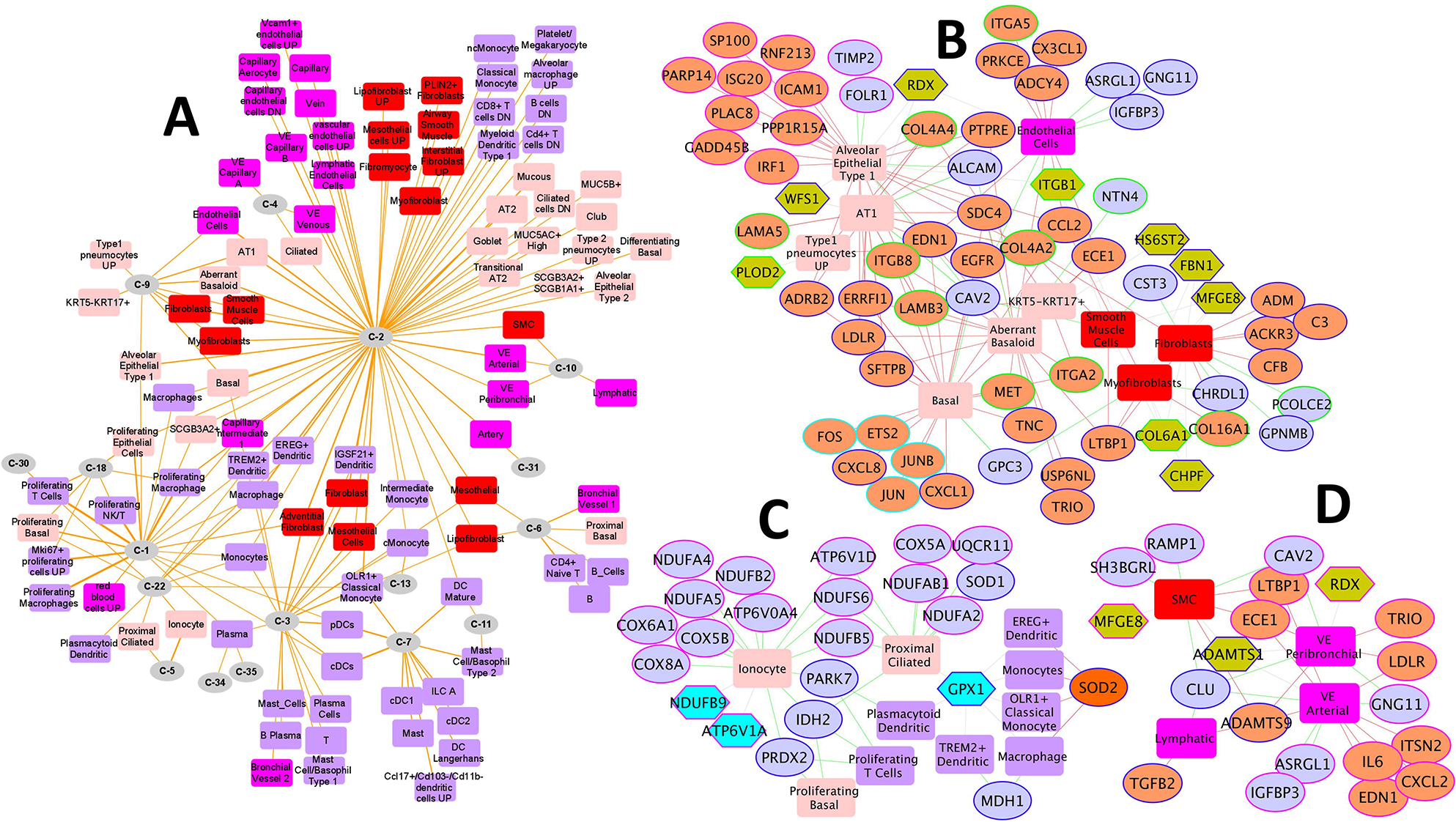
Lung single cell marker enrichments in gene clusters from the integrated interactome of conserved DEGs of SARS-CoV-2 infection models and SARS-CoV-2-host protein-protein interactions. **(A)** Marker enrichment network of 17 candidate clusters (circular nodes) which were enriched for at least one lung cell type (rectangular nodes) in normal or aging lung. **(B) – (D).** Networks of enriched cell types and the corresponding gene markers from select gene clusters. Highlighted gene clusters include C-9, C-1, C-2, and C-3 (Panel B); C-5 and C-22 (Panel C); and C-2 and C10 (Panel D). In all the networks, the rectangular nodes represent the enriched cell types and are colored based on their cell type category – immune cell types are in purple color, endothelial are in magenta, epithelial are in salmon and stromal cell types are in red color. The conserved up and downregulated genes are in orange and purple colored ellipses while the hexagon shaped nodes are genes that encode proteins interacting with SARS-CoV-2 proteome.

#### Gene clusters - Functional enrichment

To further characterize the 35 selected clusters, we next performed functional enrichment analysis against several known gene sets (Supplementary File 12). Cluster C-1 (190 genes) was enriched for innate immune response (48 genes) and type I interferon signaling (26 genes) while genes from cluster C-2 (92 genes) were involved in transport regulation (31 genes) and tube development (31 genes). We also found genes associated with abnormal cardiovascular development (21 genes) in cluster C-2. Clusters C-7 (20 genes) and C-8 (20 genes) had genes associated with abnormal interleukin and cytokine secretion phenotypes. Similarly, clusters C-12 (14 genes), C-28 (6 genes) and C-23 (8 genes) were jointly enriched for mitochondrion translation, organization, and transport. We also saw high concordance between the functional and cell type marker enrichments among the candidate clusters. Cluster C-4 (73 genes) was found to consist of markers (9 genes) in human lung ciliated cells and was simultaneously enriched for ciliary and basal body-plasma membrane docking processes. Also, cluster C-13 (14 genes) was enriched for processes associated with neutrophil activation involved in immune response and had marker genes in classical monocyte cells. Similarly, cluster C-9 (18 genes) was enriched for collagen proteins associated extracellular matrix organization (ECM) while also having several fibroblast and smooth muscle cell markers. Finally, several genes regulating circadian rhythm in mammals (*NFIL3, PER1, PER2, PER3 and SIK1*) were seen in cluster C-25 (7 genes) (Supplementary File 12).

#### Gene clusters – genotype-phenotype associations

As described in earlier sections, we repeated the enrichment for genotype-phenotype associations but this time using each of the 35 clusters. The clusters revealed enrichment for several physiological and phenotypic traits that provide several insights into the underlying pathophysiology of COVID-19 (Supplementary Files 13 and 14). For example, among the most significantly enriched traits were respiratory system disease (clusters C-7 and C-8), asthma (C-7), autoimmune disease (clusters C-7 and C-29), allergic rhinitis (C-7), immune system disease (cluster C-7 and C-8) and diabetes (C-15). We also observed risk genes associated with several inflammatory system disorders like inflammatory bowel disease and Crohn’s disease (C-7), ulcerative colitis (C-8), rheumatoid arthritis (clusters C-7 and C-8) and ankylosing spondylitis (C-8). Apart from helping in understanding the underlying pathophysiology of COVID-19, the enriched traits can potentially help the researchers to understand or formulate hypotheses surrounding the long-hauler patients or survivors. For instance, apart from being risk factors for COVID-19, could COVID-19 conversely be a risk factor for an autoimmune or neurodegenerative disease? A plausible mechanism could be through an over-activated innate immune system [70–72]. Both acute and delayed neurological and neuropsychiatric effects have been associated with previous viral pandemics [62, 63].

### Meta-analysis of results from 35 clusters and enrichment network visualization

The overall concordance between the enriched cell types and functional enrichments for different gene clusters motivated us to perform a more general analysis across all enrichments (i.e., cell type, phenotypic traits, biological processes, and pathways). To do this, we selected a subset of enriched terms (top ten enriched terms from GO-BP, Reactome pathways, mouse phenotype), cell types, and traits (both PheGenI and GWAS Catalog) from each of the 35 clusters, and converted them into a network layout. Gephi (https://gephi.org), an open-source graph visualization platform [73], was used to construct and visualize the network. Enriched terms (e.g., biological processes, pathways, phenotypic traits, cell types) are represented as nodes and two nodes are connected if they share at least one or more of the 35 gene clusters from the DEG and SARS-CoV-2-host interactome map. This resulted in a dense network of 1213 nodes and 32,455 edges (Supplementary File 15). The substructure of this network was investigated by estimating community membership modules using the Louvain algorithm (implemented in Gephi) [74]. With a resolution set to 0.3, we found 31 communities of highly interconnected biological processes, pathways, cell types, and phenotypic traits and a high modularity score of 0.676 (Figure 6). Since sub-units of a functional complex (a cluster of pathways, biological processes, phenotype, etc.) work towards the same biological goal, prediction of an unknown pathway or biological process or a phenotype as part of this complex also allows increased confidence in the annotation of that functional cluster. Additionally, by doing this, potential redundancies across different sources (e.g., ontology or cell types) could be reduced, apart from enabling interpretation of the enrichment results through intra-cluster and inter-cluster similarities of enriched terms [75]. For instance, cluster C-9 (18 genes with 5 genes encoding proteins that directly interact with SARS-CoV-2 and 13 genes that are conserved DEGs in SARS-CoV-2 infection models) shows enrichment of several pathways, biological processes and cell types that are semantically related. Among the enriched biological processes and pathways are cell substrate adhesion, ECM, collagen metabolism while the cell type enrichment showed myofibroblasts, smooth muscle cells and fibroblasts (Figure 6). Similarly, clusters C-7 and C-8 were enriched for toll-like receptor signaling pathway, cytokine-cytokine receptor interaction, NF-KappaB signaling, CD40 signaling. These clusters showed enrichment for abnormal interleukin secretion and T-cell physiology and for several GWA loci such as granulocyte count, inflammatory biomarker measurement, Crohn’s disease, and ulcerative colitis (Figure 6).

**Figure 6:**
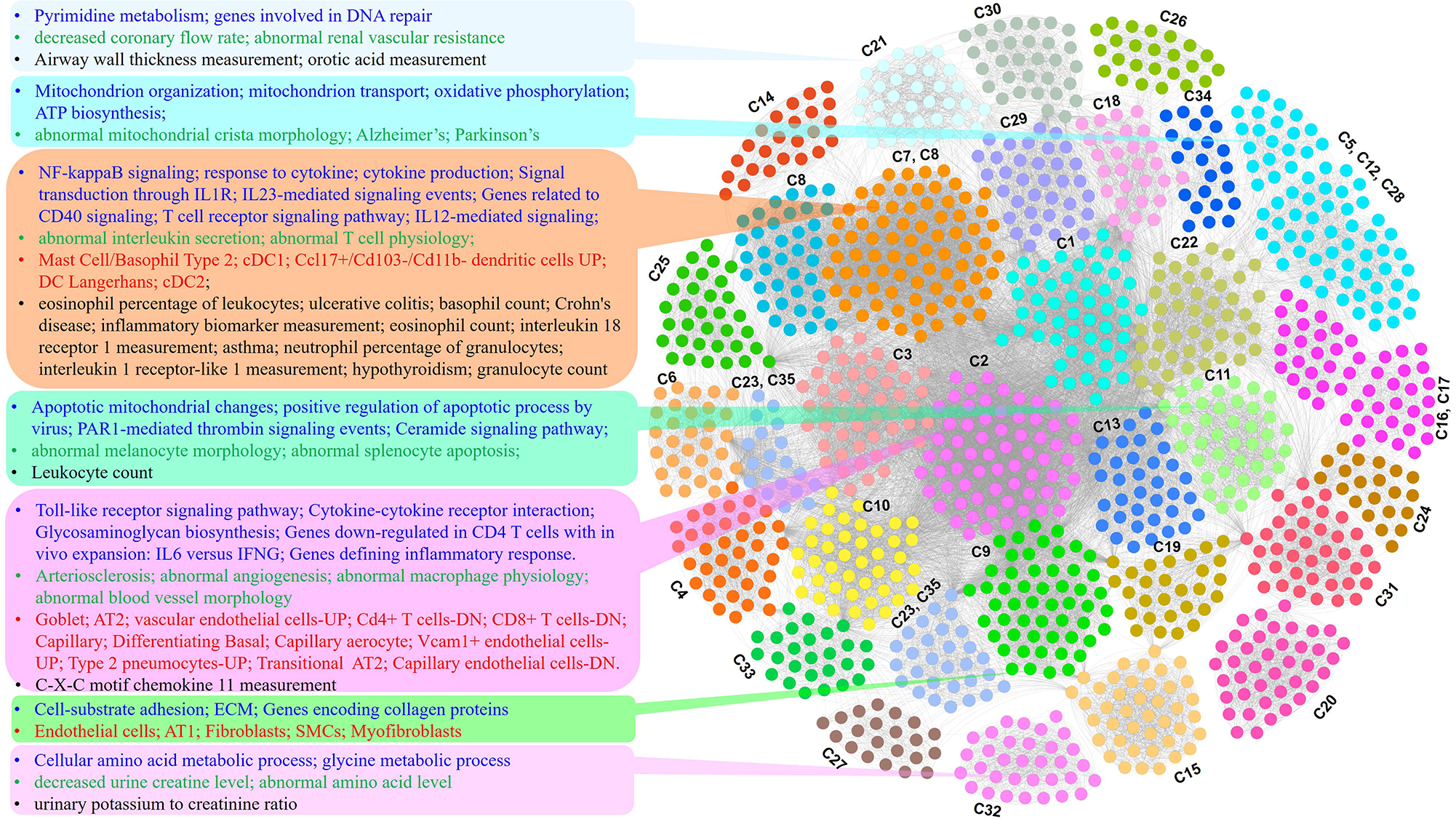
Network visualization of the results from joint analysis of multiple annotations from the 35 gene clusters. Network representation of clustered enriched terms from functional enrichment analysis (multiple annotations such as gene ontology, pathways, lung cell types, and GWA loci) of candidate gene clusters from the integrated conserved DEG and SARS-CoV-2-host interacting protein network. Enriched terms from different annotation categories are represented as nodes while the edges represented shared gene clusters (35 select gene clusters). Representative terms for some of the clustered enrichment term modules are listed on the left side with different font colors representing different annotation categories (blue-biological processes/pathways; green-phenotype; red, cell type; black-GWA trait) of the font. The underlying gene clusters (35 select gene clusters) for each of the clustered functional terms are also shown.

## DISCUSSION

Although the body of knowledge surrounding COVID-19/SARS-CoV-2 is recent, it is growing at a rapid pace. Currently, there are no antiviral treatments nor vaccines against this disease and the long-term consequences in recovered patients is not known. Nevertheless, leveraging the accumulating COVID-19/SARS-CoV-2 omics data, extrapolations can be drawn and hypotheses generated about the potential effects of the virus on different cell types, physiological functions, and distinct features or pathways with the premise that all of these can be exploited for potential drug discovery. By integrating and analyzing the SARS-CoV-2 transcriptomic data in the context of SARS-CoV-2-human virus-host protein interaction map, single cell signatures of normal and aging lung, gene annotations, and human genotype-phenotype associations, we have identified several functional modules that can have direct bearing on furthering the understanding of this devastating pandemic.

The various categories of cellular functions and phenotypic traits found by meta-analyses included both expected and novel biological insights of COVID-19. For instance, metanalysis of the transcriptome revealed a strong enrichment for several neurologic - physiological and pathological - traits. Interestingly, about 9% of patients with COVID-19 are reported to develop confusion or dizziness, and some are reported to have anosmia and ageusia [68, 76–78]. Although it is not known whether SARS-CoV-2 can directly infect the central nervous system, the fact that another beta coronavirus (HCoV-OC43) has been associated with fatal encephalitis [69] necessitates a closer scrutiny of the neurological implications – acute and chronic - of COVID-19.

Our study holds certain limitations. The transcriptomics data used is from in vitro and in vivo model systems with no samples from human patients of COVID-19. Although there are emerging data sets from COVID-19 patients, they are currently limited. For example, from the study that reported SARS-CoV-2-triggered transcriptome in Calu-3 cells, transcriptomics data from lung samples of COVID-19 patient are available. However, these samples were from just one patient. Nevertheless, the conserved signature from our study can be used to compare with more robust transcriptomic signatures from COVID-19 patients as and when they are available. For example, comparing the DEGs from nasopharyngeal swabs from human patients (GSE152075) with the DEGs from the three SARS-CoV-2 infection models showed strong concordance with the Calu-3 and the humanized mouse model (Supplementary File 16). Surprisingly, there was no significant correlation between the VeroE6 upregulated genes and COVID-19 patient nasopharyngeal upregulated genes. Although Vero E6 is the most widely used cell line to replicate and isolate SARS-CoV-2 [4], the expression level of TMPRSS2, the receptor that SARS-CoV-2 uses to prime the spike protein of SARS-CoV-2 [80, 81] is reported to be quite low in this clone. Additionally, there were many DEGs (1180 upregulated and 1734 downregulated) that are specific to human patients suggesting the inherent limitations of current in vitro and in vivo models of COVID-19.

The STRING-based PPI network data, as is the case with any omics data, suffers from data incompleteness and certain degree of noise. Further, there are no clear guidelines on what to use for the STRING interaction score cutoff. Similarly, although Markov clustering is recommended for module detection [79], there exist no guidelines for inflation factor threshold nor for the functional annotation of modules. Nevertheless, to overcome some of these limitations, we used a very stringent cut-off score of 0.9 for STRING interactions. Similarly, we selected 2.5 (default) as the MCL inflation factor which generated 153 clusters of 2–190 genes. The cluster composition can vary depending on the clustering algorithm and parameters. For enrichment analysis we used 35 of those clusters that have 5 or more genes. Lastly, our study does not include any experiment validations. However, the gene clusters, enriched biological processes and pathways, the normal and aging lung marker and human genotype-phenotype associations emerging from our joint analysis of COVID-19 and non-COVID-19 related data can serve as valuable resource for the scientific community to formulate or further investigate hypotheses. However, we caution against interpreting our results from genotype-phenotype enrichment analysis as causal effects. Although we found enrichment for smoking and COPD, blood pressure, and body mass index (along with several other traits), as a recent study pointed out [56], carefully designed causal analyses focusing on the causal effects of enriched traits and COVID-19 outcomes are needed.

In conclusion, with no therapeutic prevention or intervention methods available, preclinical research using in vitro and in vivo model organisms is needed to understand SARS-CoV-2 infection, clinical manifestations of COVID-19, and to test therapeutic and preventive agents for safety and efficacy. Based on current in vitro and in vivo models of SARS-CoV-2 infection, we believe that our conserved gene signature from SARS-CoV-2 infection models, SARS-CoV-2 targeted human protein clusters, and their downstream analyses will help researchers to further understand and formulate testable hypotheses for COVID-19 risk factors, pathophysiology, potential sequelae, and support development of therapeutic agents.

## Ethics approval and consent to participate

Not applicable.

## Consent for publication

Not applicable.

## Availability of data and materials

All data generated or analyzed during this study are included in this published article and its supplementary information files.

## Author contributions

S.G. and A.J. conceived this study. S.G., M.S. and A.J. collected and analyzed data. S.G. and A.J. interpreted results from data. S.G., M.S. and A.J. edited this manuscript. S.G. and A.J. wrote the first draft.

## Acknowledgements

Not applicable.

## Supplementary Data

**Supplementary File 1: SARS_CoV2_DEGs:** COVID-19 DEGs from human (Calu-3) and non-human primate (VeroE6) cell lines and from a mouse model (Ad5-hACE2).

**Supplementary File 2**: **Aging markers in lung and liver**: DEGs in aging lung (50-79 years vs 20-29 years) and liver (50-69 years vs 20-29 years) tissues from GTEx database. Human aging associated genes from GenAge are also included.

**Supplementary File 3**: **Conserved SARS-CoV-2 DEGs:** Conserved DEGs (833 upregulated and 634 downregulated) which are differentially regulated in at least 2 out of the 3 studies (human Calu-3, non-human primate (VeroE6) cell lines and a mouse model (Ad5-hACE2). Also included is the “core” set of 147 DEGs (106 upregulated and 41 downregulated) dysregulated in all the three studies.

**Supplementary File 4: Functional terms enriched in core SARS-CoV-2 signature:** Results from functional enrichment analysis of core SARS-CoV-2 signature (separately for 106 upregulated and 41 downregulated genes) using the ToppFun application of the ToppGene Suite.

**Supplementary File 5: Functional terms enriched in SARS-CoV-2 conserved signature:** Results from functional enrichment analysis of conserved SARS-CoV-2 signature (separately for 833 upregulated and 634 downregulated genes) using ToppFun application.

**Supplementary File 6: Elsevier pathways enriched in SARS-CoV-2 conserved signature:** List of enriched Elsevier pathways enriched among the conserved SARS-CoV-2 signature (separately for 833 upregulated and 634 downregulated genes). Pathway Collection available as part of the Enrichr application.

**Supplementary File 7**: **Cell marker enrichments in SARS-CoV-2 conserved signature:** Results from enrichment analysis of the SARS-CoV-2 conserved DEGs against single cell marker gene sets from three different human lung scRNA-seq studies and also aging lung cell markers in mice.

**Supplementary File 8: Phenotype-Genotype Integrator (PheGenI) traits enriched in SARS-CoV-2 conserved DEGs:** Both physiological traits and also the human disease loci from NCBI PheGenI (https://www.ncbi.nlm.nih.gov/gap/phegeni) enriched in conserved SARS-CoV-2 signature.

**Supplementary File 9: Genome-wide association (GWA) traits from GWAS Catalog enriched in SARS-CoV-2 signature:** GWA traits from GWAS Catalog (https://www.ebi.ac.uk/gwas/downloads) enriched among the SARS-CoV-2 DEGs. Associations of the child traits, parsed from the experimental factor ontology (EFO) hierarchy, were also used.

**Supplementary File 10: SARS-CoV-2 conserved DEG and human interaction map:** SARS-CoV-2 human interaction map constructed from a joint interactome of SARS-CoV-2 conserved signature and SARS-CoV-2-human interaction map. We found 153 clusters in total and found 35 candidate clusters with a minimum of 5 genes.

**Supplementary File 11: Cell marker enrichments in SARS-CoV-2 human interaction map clusters:** Enriched cell types from three different human lung scRNA-seq studies and aging lung cells in mice within the 35 candidate gene clusters.

**Supplementary File 12: Results from functional enrichment analysis in SARS-CoV-2 human interaction map clusters:** Enriched functional terms (Gene Ontology biological processes, mouse phenotypes, biological pathways from Reactome and co-expressed genesets) among the 35 candidate clusters from the SARS-CoV-2 human interaction map.

**Supplementary File 13: Phenotype-Genotype Integrator (PheGenI) traits enriched in SARS-CoV-2 human interaction map clusters:** Both physiological and phenotypic traits from NCBI PheGenI enriched among the SARS-CoV-2 human interaction map clusters.

**Supplementary File 14**: **Genome-wide association (GWA) traits from GWAS Catalog enriched in SARS-CoV-2 human interaction map clusters:** GWA traits from GWAS Catalog enriched among the SARS-CoV-2 human interaction map clusters. Associations of the child traits were also included.

**Supplementary File 15: Meta-analysis network based on enriched cell types and functional enrichments from candidate SARS-CoV-2 human interaction map clusters:** A Meta-analysis substructure where nodes are enriched terms while links are the clusters shared. The final network comprised 1213 nodes and 32,455 edges.

**Supplementary File 16**: Comparison of DEGs from SARS-CoV-2 infection models with that of DEGs from human nasopharyngeal swabs of COVID-19 patients.

