## Supplementary figures and images for "Secondary analysis of transcriptomes of SARS-CoV-2 infection models to characterize COVID-19"

### Supplementary File 16

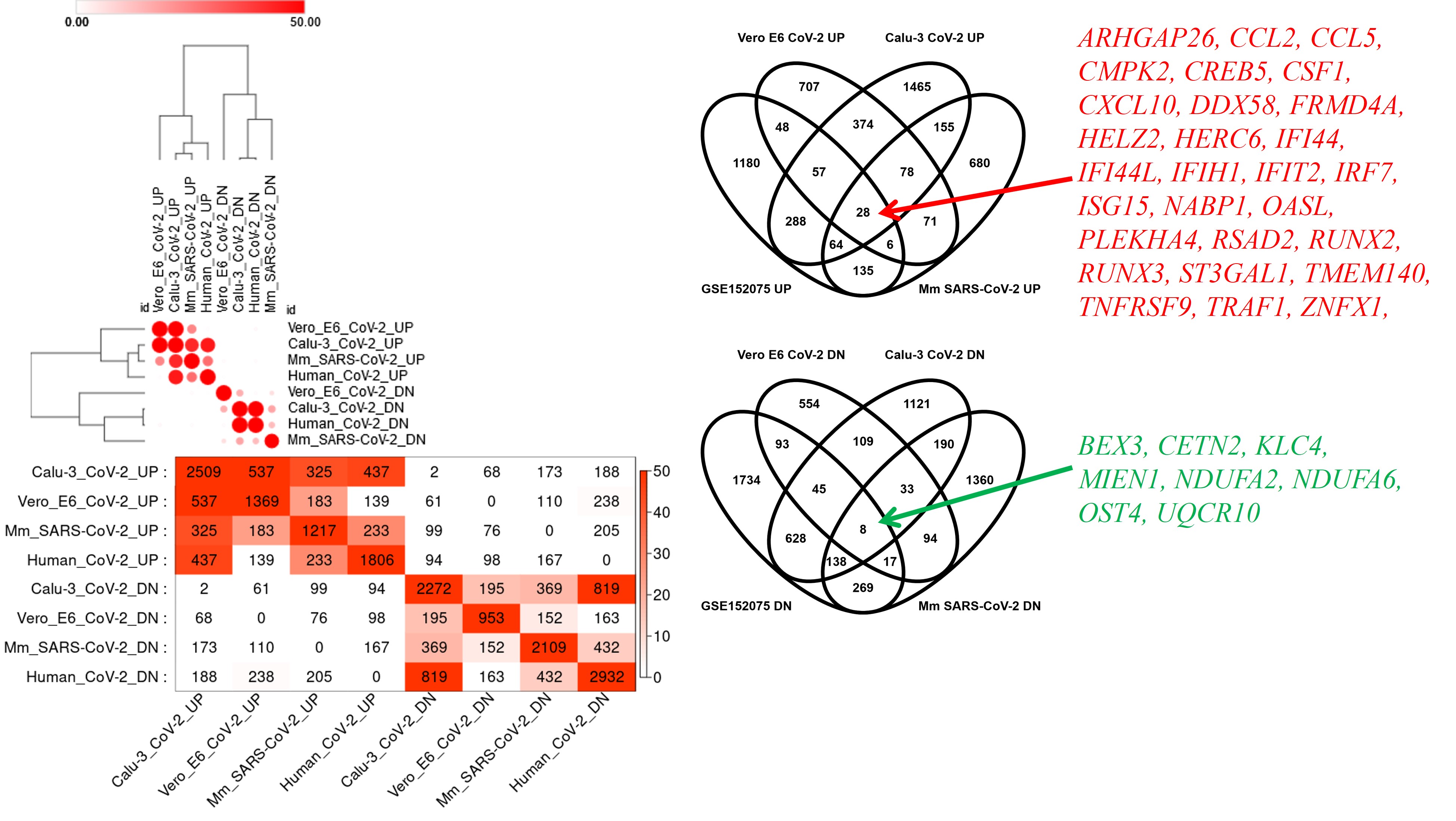
